# Comparing sequence and structure of falcipains and human homologs at prodomain and catalytic active site for malarial peptide-based inhibitor design

**DOI:** 10.1101/381566

**Authors:** Thommas M. Musyoka, Joyce N. Njuguna, Özlem Tastan Bishop

**Author notes:** Corresponding author details: Research Unit in Bioinformatics (RUBi), Department of Biochemistry and Microbiology, Rhodes University, P.O. Box 94, Grahamstown, 6140, South Africa.

## Abstract

Falcipains are major cysteine proteases of *Plasmodium falciparum* essential in hemoglobin digestion. Several inhibitors blocking their activity have been identified, yet none of them has been approved for malaria treatment. For selective therapeutic targeting of these plasmodial proteases, identification of sequence and structure differences with homologous human cathepsins is necessary. The protein substrate processing activity of these proteases is tightly controlled in space and time via a prodomain segment occluding the active site making it inaccessible. Here, we utilised *in silico* approaches to determine sequence and structure variations between the prodomain regions of plasmodial proteins and human cathepsins. Hot spot residues, key for maintaining structural integrity of the prodomains as well as conferring their inhibitory activity, were identified via residue interaction analysis. Information gathered was used to design short peptides able to mimic the prodomain activity on plasmodial proteases whilst showing selectivity on human cathepsins. Inhibitory potency was highly dependent on peptide amino acid composition and length. Our current results show that despite the conserved structural and catalytic mechanism of human cathepsins and plasmodial proteases, significant differences between the two groups exist and may be valuable in the development of novel antimalarial peptide inhibitors.

## Introduction

Malaria, caused by parasites from the genus *Plasmodium* and transmitted to human by a female anopheles mosquito bite, is still a devastating disease even though the global incidences have drastically dropped in recent years [1]. Parallel to evolving mosquito resistant to insecticides [1–4], continuously emerging resistant strains of parasite to current drugs [5–8] present an immense challenge for the eradication of malaria. A recent study promisingly showed that pre-existing resistance may not be a major problem for novel-target antimalarial candidates, and fast-killing compounds may result in a slower onset of clinical resistance [9]. Hence, the identification and development of alternative anti-malarial inhibitors with novel mode of action against new as well as known drug targets with certain key features are very important.

Proteases are considered as good parasitic drug targets and details are presented in a number of articles [10–16]. Cysteine proteases have a central role in *Plasmodium* parasites during hemoglobin degradation [17,18], tissue and cellular invasion [19], activation of pro-enzymes [20,21], immunoevasion and egression [11,21,22]. Red blood cell (RBC) invasion and rupturing processes as well as intermediate events involving hemoglobin metabolism are characterised by increased proteolytic activity. During the asexual intraerythrocytic stage, *Plasmodium* parasites degrade nearly 75% of host RBC hemoglobin [23,24] to acquire nutrients as they lack a *de novo* amino acid biosynthetic pathway. By this process, they can acquire all their amino acid requirements necessary for growth and multiplication with an exception of isoleucine which is exogenously imported as it is absent in human hemoglobin [10,25,26]. Hemoglobin degradation is an intricate and efficient multistage protein catabolic process occurring inside the acidic food vacuole [18,27].

This study focuses on a subgroup of papain-like Clan CA plasmodial cysteine proteases, namely Falcipains (FPs) of *P. falciparum* and their homologs. *P. falciparum* has four FPs; FP-1, FP-2, FP-2’ and FP-3. FP-1 is the most conserved protease among the four proteases, and its role in parasite entry into RBCs is yet to be resolved. Although its inhibition using specific peptidyl epoxides blocked erythrocyte invasion by merozoites [28], FP-1 gene disruption in blood stage parasites does not affect their growth [29,30]. Despite its biological function remaining uncertain, FP-2’ is biochemically similar to FP-2 and shares 99% sequence identity [22,31]. FP-2 (FP-2’) and FP-3 share 68% sequence identity and are the major cysteine proteases involved in hemoglobin degradation in the parasite [32–35]. Expression of these proteins during the blood stage by plasmodia is strictly regulated in a site-specific and time-dependent manner [28,36,37]. These hemoglobinases have differential expression timing during the trophozoite stage: the early phase is characterised by FP-2 abundance while FP-3 is abundant at the late stages [17,22]. It was shown that targeted disruption of FP-2 gene in plasmodia results in accumulation of undigested hemoglobin in the food vacuole and its enlargement [17], therefore the protein can be considered as a promising drug target [38,39]. On the other hand, inhibiting individual proteases might not be essential due to redundancy in the hemoglobin digestion stage [10], hence any inhibitor design for FPs should consider blocking the activity of both FP-2 and FP-3. The importance of FP-2 as a drug target was also indicated in a recent study in which FP-2 polymorphisms were shown that are associated with artemisinin resistance [40].

Other *Plasmodium* species also express proteins highly homologous to FP-2 and FP-3 [41–44]. These include vivapains (vivapain 2 [VP-2] and vivapain 3 [VP-3]), knowlesipains (knowlesipain 2 [KP-2] and knowlesipain 3 [KP-3]), berghepain 2 [BP-2], chabaupain 2 [CP-2] and yoelipain 2 [YP-2] from *P. vivax, P. knowlesi, P. berghei, P. chabaudi* and *P. yoelii* respectively. All these proteins are related both in sequence and function to the papain-like class of enzymes including human cathepsins. The plasmodial proteases have, however, unusual features compared to the human ones including, much longer prodomains and specific inserts in the catalytic domain - a “nose” (~ 17 amino acids) and an “arm” (~ 14 amino acids) [37,45,46]. In native environment, cysteine proteases are regulated either by their prodomain (zymogen form) or by other endogenous macromolecules like cystatins [47,48] and chagasin [49]. During erythrocyte entry, *P. falciparum* parasites secrete falstatin, a potent picomolar inhibitor of both FP-2 and FP-3 thus regulating the activity of these proteases on important surface proteins required for invasion [19,48]. In the zymogen form (Figure 1), a part of the prodomain flips over the active pocket and its subsites located on the catalytic domain [50], blocking its enzyme activity [51]. The acidic environment within a food vacuole (plasmodia) or lysosome (humans) triggers prodomain cleavage thus activating the catalytic domain [52,53].

**Figure 1.**
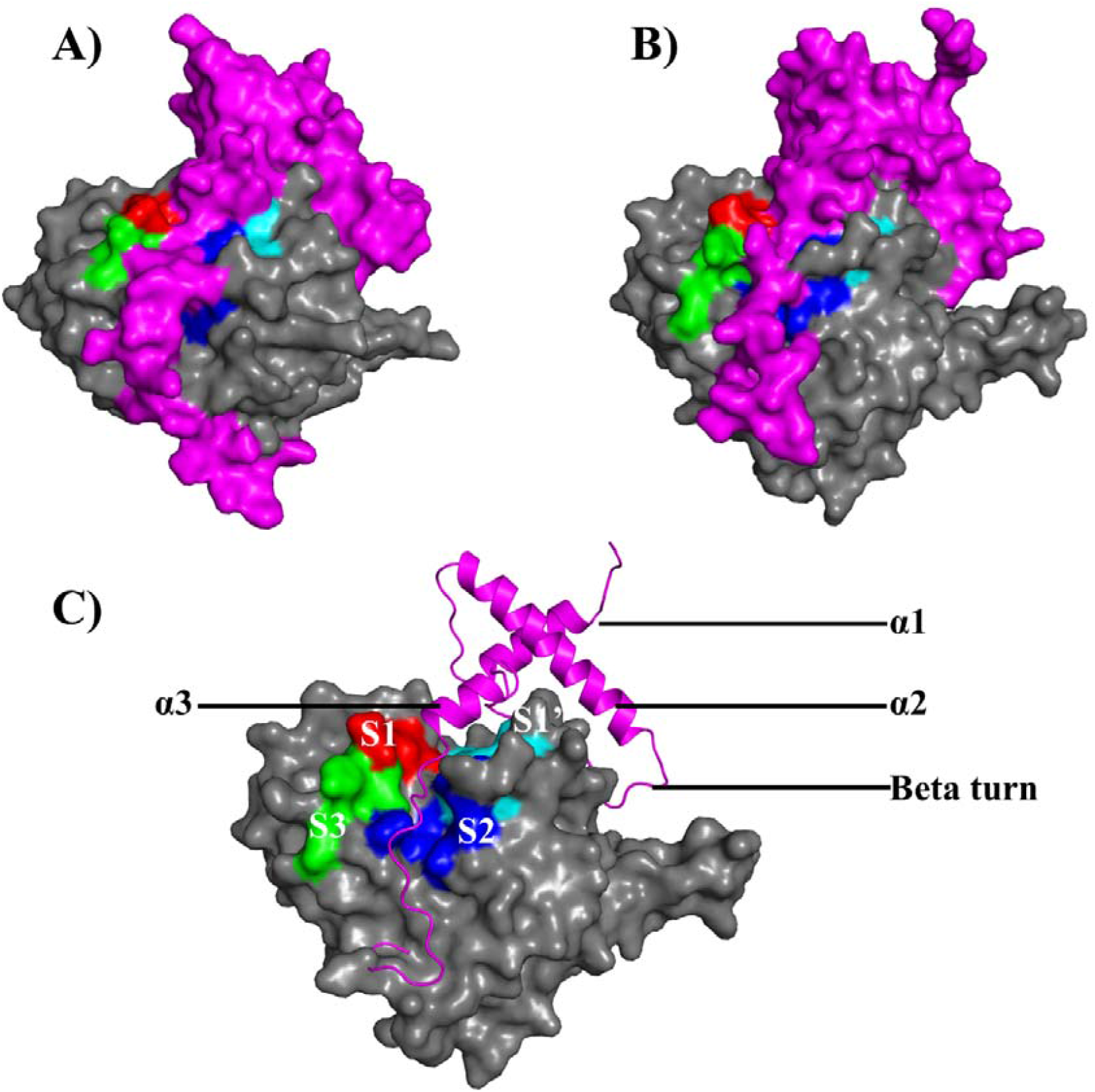
Clan CA cysteine protease zymogen prodomain-catalytic domain interaction modes. Surface representation of A) human Cat-K and B) FP-2. C) FP-2 prodomain structural elements (pink; in cartoon representation) interacting with the S1 (red), S2 (blue), S3 (green) and S1’ (cyan) subsites of the catalytic domain.

The literature comprises a large number of inhibitor studies against FPs including peptide-based [31,54–56], non-peptidic [50,57–61] and peptidomimetic [58,62,63] studies. Hitherto, none of these inhibitors has been approved as an antimalarial drug as they have limited selectivity against host cathepsins, homologs to the parasites proteases. To overcome this, distinctive features between these two classes of proteins must be determined. Primarily, the current work utilises *in silico* approaches to characterize FP-2 and FP-3, their homologs from other *Plasmodium* species as well as human homologs (cathepsins) to identify sequence, physicochemical and structure differences that can be exploited for peptide-based antimalarial drug development. Although the two protein classes share high similarity, important differences that can be essential for inhibitor selectivity exist [50,64]. Our main aim in this study is to elucidate the inhibitory mechanism of plasmodial prodomain region responsible for endogenous regulation of the catalytic domain, information which may be useful in the design of novel inhibitors. For this purpose, using domain-domain interaction approaches, specific hot spot residues critical for the mediation of the prodomain inhibitory effect were identified.

To further identify a potential peptide segment, which could strongly bind to the plasmodial catalytic domains and mimic the native prodomain inhibitory effect, five short peptide sequences based on the identified hot spot residues were suggested. Flexible docking of these peptides against the catalytic domains identified a short 13-mer oligopeptide with preferential binding towards plasmodial proteases. This oligopeptide could be a starting platform for the development and testing of novel peptide based antimalarial therapies against plasmodial cysteine proteases.

## Material and methods

A workflow consisting of the different methods, tools and databases used in this study is shown in Figure 2. Unless otherwise indicated, amino acid numbering is based on individual protein full length as listed in Table S1.

**Figure 2.**
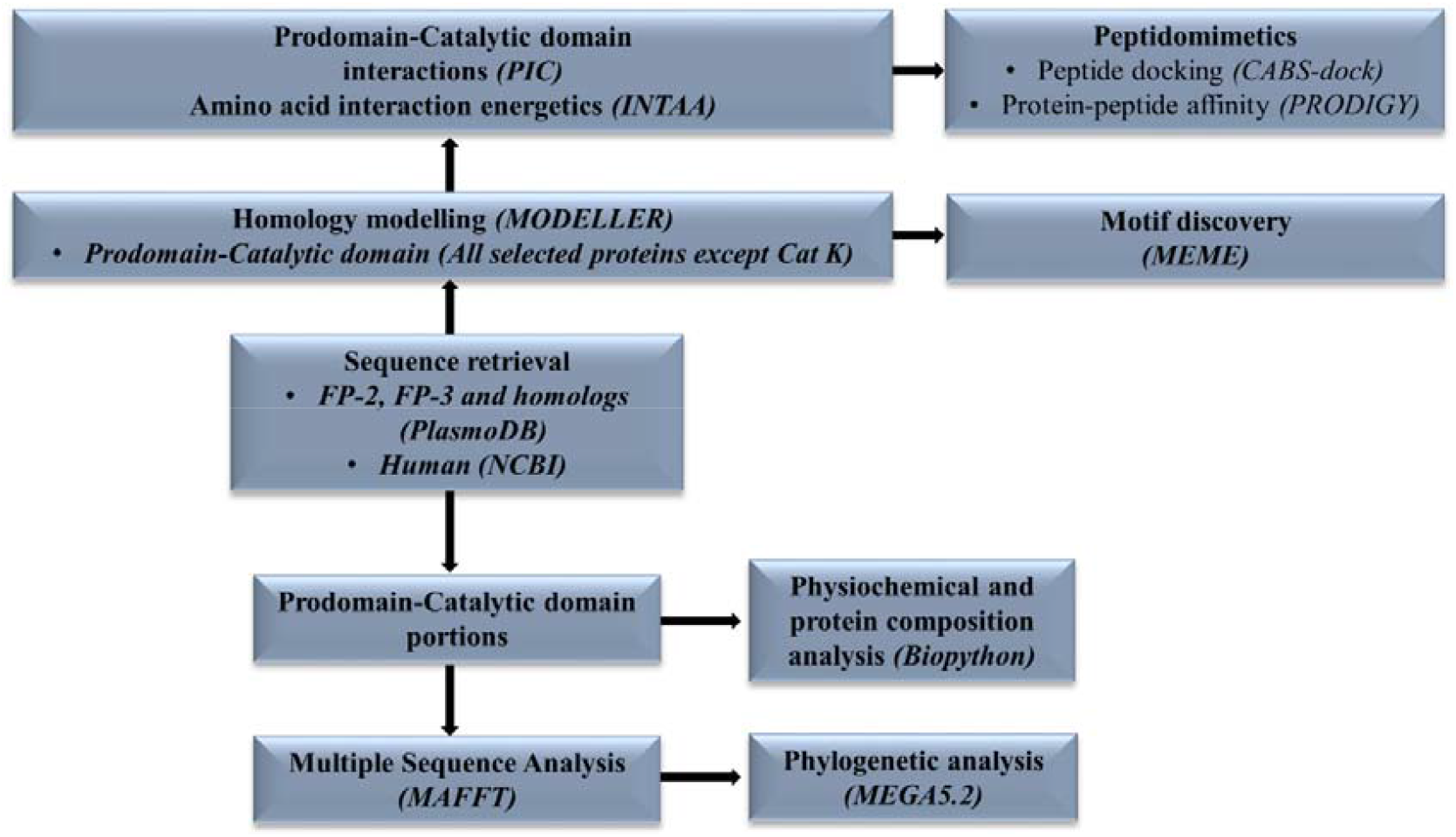
A graphical workflow of the methods and tools (in brackets) used in sequence and structural analysis of FP-2, FP-3 and their homologs.

### Sequence retrieval and multiple sequence alignment

Using FP-2 (PF3D7_1115700) and FP-3 (PF3D7_1115400) as query sequences, seven plasmodial protein homologs together with three human homologs (Table 1) were retrieved from the PlasmoDB version 9.31 [65] and NCBI [66] databases respectively as described earlier [50]. A pronounced feature present in the cathepsin L (Cat-L) like plasmodial proteases is the presence of an N-terminal signalling (non-structural) peptide sequence (~150 amino acids), which is responsible for targeting them into the food vacuole. For each of the plasmodial proteins, this segment was chopped off, and the remaining prodomain-catalytic portion saved into a Fasta file (Text S1). As guided by the partial zymogen complex crystal structure of Cat-K [PDB: 1BY8], ~ 21 amino acids (N-terminal) were also chopped off from the human cathepsin prodomain sequences. Together, these sequences were used in the rest of the study, and are referred as “partial zymogen” or “prodomain-catalytic domain” sequences interchangeably in the manuscript. Position details of the prodomain and catalytic portions per protein are listed in Table S1. To determine the conservation of the prodomain-catalytic portion, multiple sequence alignment (MSA) was performed using PROfile Multiple Alignment with predicted Local Structures and 3D constraints (PROMALS3D) web server [67] with default parameters except PSI-BLAST Expect value which was adjusted to 0.0001, and the alignment output visualised using JalView [68].

**Table 1.**
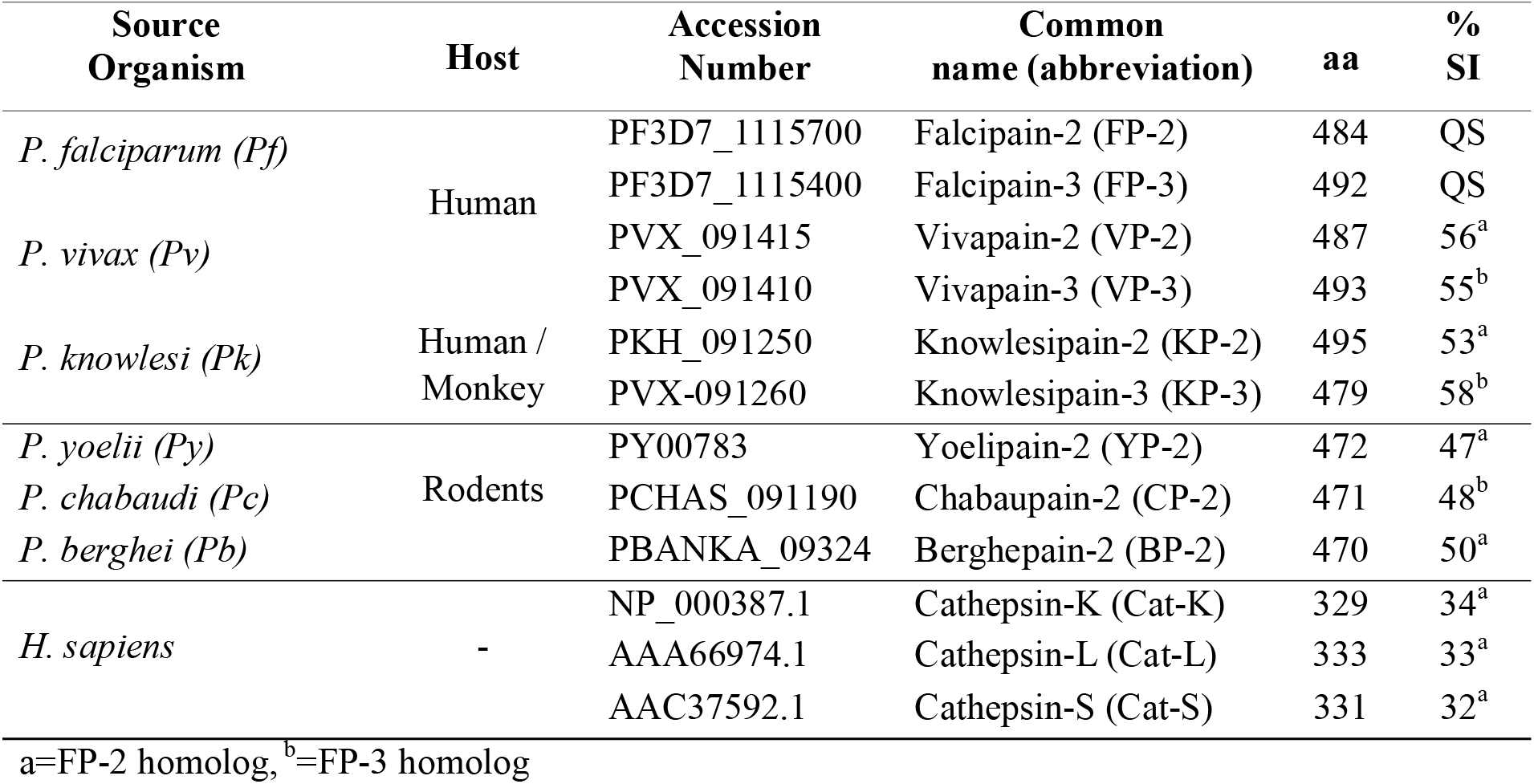
Details of all protein sequences retrieved from PlasmoDB and NCBI databases. Percentage sequence identity (SI) is calculated based on the partial zymogen of query sequence (QS) and that of corresponding homolog.

### Phylogenetic inference

Using Molecular Evolutionary Genetic Analysis (MEGA) version 5.2 software [69], the evolutionary relationship of plasmodial proteases and human cathepsins was evaluated with the following preferences; Maximum Likelihood (statistical method) and Nearest-Neighbor-Interchange (NNI) as the tree inference option. A total of 48 amino acid substitution models were calculated for both complete (100%) and partial (95%) deletion and the best three models based on Bayesian Information Criterion (BIC) were selected (Table S2). For each selected model, the corresponding gamma (G) evolutionary distance correction value was selected to build different phylogenetic trees and comparison was made to determine robustness of dendrogram construction process. *Toxoplasma gondii* Cat-L [NCBI accession number: ABY58967.1] was included in the tree calculations as outgroup.

### Physicochemical properties

Using an *ad hoc* Python and Biopython script, the amino acid composition and physicochemical properties, namely molecular weight (Mr), isoelectric point (pI), aromaticity, instability index, aliphatic index and grand average of hydropathy index (GRAVY) of the proteins were determined.

### Motif analysis

Multiple Em for Motif Elicitation (MEME) standalone suite version 4.10.2 [70] was used to identify the composition and distribution of protein motifs within partial zymogen sequences. A Fasta file (Text S1) containing sequence information of the different proteins was parsed to MEME software with analysis preferences set as; -nostatus –time 18000 –maxsize 16000 – mod zoops –nmotifs X –minw 6 –maxw 50. The variable X (a whole number from 1) was varied until no more unique motifs were assessable as determined by motif alignment search tool (MAST) [71]. A heat map showing motif distribution was generated using an in house Python script. PyMOL was used to map the different motifs onto the protein structures (The PyMOL Molecular Graphics System, Version 1.6.0.0 Schrödinger, LLC).

### Homology modelling and structure validation

MODELLER version 9.18 [72] was used to build homology models of the inhibitor complex of all proteins except for Cat-K which has already a crystal structure. Using a combination of templates, high quality prodomain-catalytic domain complexes of the plasmodial proteases as well as cathepsins (Cat-L and Cat-S) were calculated by MODELLER with refinement set to very slow. Table S3 shows the details of templates selected for each protein model. For the plasmodial proteases, the crystallographic structure of procathepsin L1 from *Fasciola hepatica* [PDB: 2O6X] was used as it had the highest similarity with most target sequences (30-38%) and high resolution of 1.40 Å. However, it lacked the arm (β-hairpin) region while the nose residues were missing. To overcome these challenges, Cat-K [PDB: 1BY8] together with FP-2 [PDB: 2OUL] (for FP-2, VP-2, KP-2, BP-2 and YP-2) and FP-3 [3BWK] (for FP-3, VP-3, KP-3 and CP-2) were additionally used. For Cat-L and Cat-S, only two templates were used [PDB: 1BY8 and 2O6X]. For each protein, 100 models were calculated and ranked according to normalized discrete optimized protein energy (Z-DOPE) score [73]. The top three models per protein were further validated using ProSA [74], Verify3D [75], QMEAN [76] and PROCHECK [77] and the best quality model selected. Table 2 shows the quality scores obtained from different homology modelling assessment tools for the top model per protein. All validation methods gave consistently high quality scores for selected models and thus could be used for further experiments. QMEAN results showed that only small portions of the loop regions in Cat-L, Cat-S, and CP-2 were built with poor quality, while the majority of the prodomain-catalytic core region in all of the proteins was accurate (Figure 3 and Figure S1).

**Figure 3.**
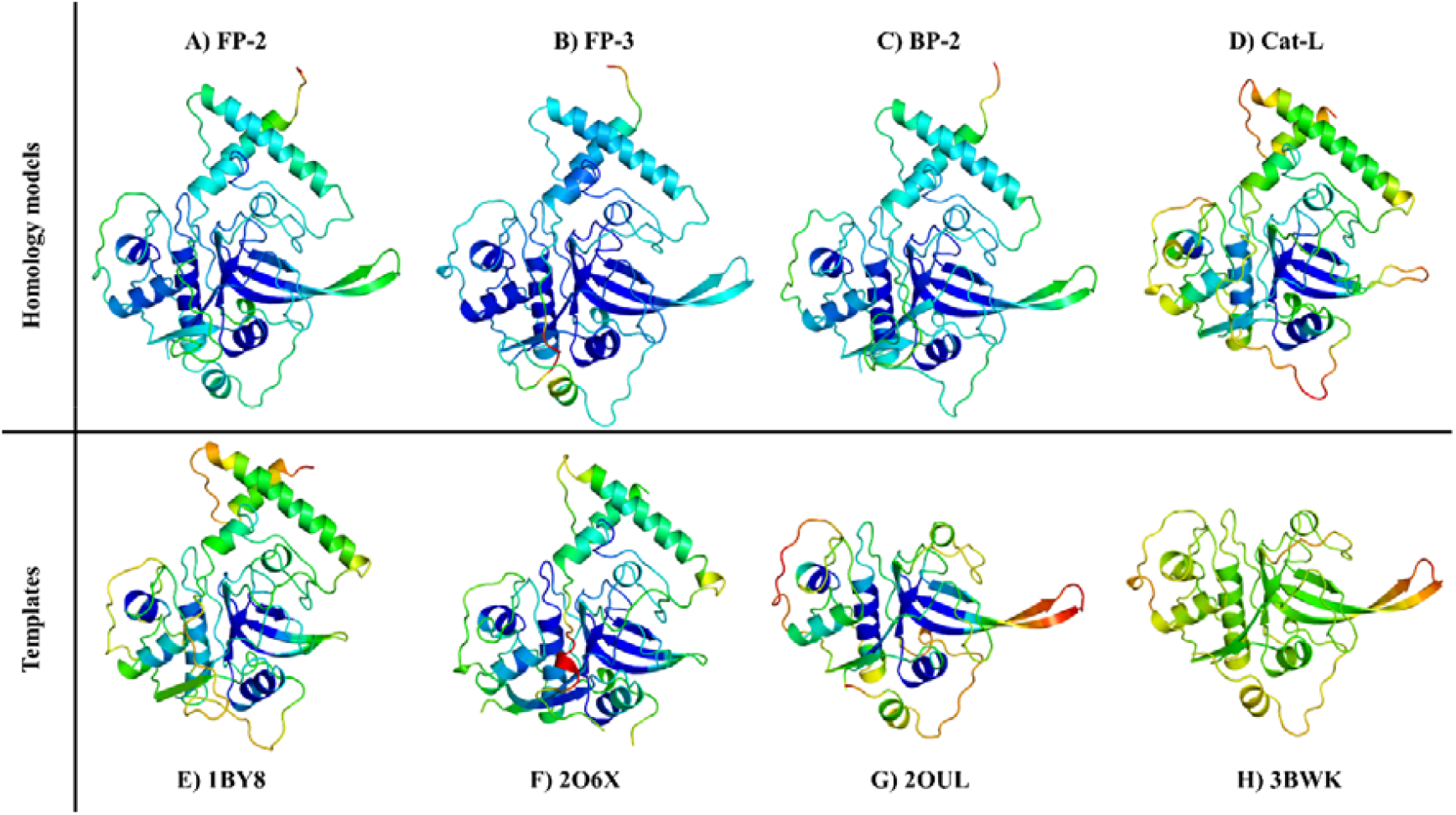
Homology models of different plasmodial proteases and human Cat-L together with the templates used in homology modelling. Colour code ranging from blue (accurate modelling) to red (poorly modelled regions).

**Table 2.** Homology model quality validation scores of partial zymogen complexes using different assessment tools.

| Protein | Z-DOPE | Verify3D | ProSA | QMEAN | RAMACHADRAN (%) |  |  |
| --- | --- | --- | --- | --- | --- | --- | --- |
|  |  |  |  |  | Favoured | Allowed | Disallowed |
| **FP-2** | -1.05 | 78.48 | -8.27 | 0.69 | 88.90 | 10.50 | 0.60 |
| **FP-3** | -0.94 | 84.64 | -7.84 | 0.67 | 89.70 | 10.30 | 0.00 |
| **VP-2** | -0.64 | 81.27 | -6.94 | 0.62 | 85.40 | 13.90 | 0.70 |
| **VP-3** | -0.74 | 79.58 | -7.31 | 0.70 | 88.60 | 11.40 | 0.00 |
| **KP-2** | -0.63 | 85.89 | -7.23 | 0.62 | 89.30 | 9.70 | 1.00 |
| **KP-3** | -0.92 | 90.63 | -7.88 | 0.63 | 86.10 | 13.90 | 0.00 |
| **BP-2** | -0.62 | 86.85 | -7.75 | 0.63 | 86.60 | 12.40 | 1.00 |
| **CP-2** | -0.60 | 84.45 | -7.02 | 0.63 | 83.90 | 13.80 | 2.30 |
| **YP-2** | -0.42 | 75.54 | -7.28 | 0.62 | 84.90 | 14.40 | 0.70 |
| **Cat-L** | -1.47 | 87.38 | -7.94 | 0.87 | 89.80 | 9.80 | 0.40 |
| **Cat-S** | -1.61 | 85.39 | -8.57 | 0.79 | 89.20 | 10.40 | 0.20 |
| **1BY8** | * | 85.16 | -8.62 | 0.78 | 65.80 | 34.20 | 0.00 |
| **2O6X** | * | 94.77 | -7.00 | 0.77 | 90.50 | 9.50 | 0.00 |
| **2OUL** | * | 98.13 | -7.90 | 0.75 | 88.10 | 11.20 | 0.70 |
| **3BWK** | * | 93.51 | -7.35 | 0.65 | 86.10 | 13.30 | 0.60 |

\*=Template

As these loop regions were far from the catalytic pocket, the resulting models were considered acceptable for further analysis.

### Prodomain-catalytic domain interaction studies and short inhibitor peptide design

To determine the prodomain inhibitory mechanism, residue interactions between prodomain and catalytic domain of plasmodial and human partial zymogen complexes were evaluated using the Protein Interaction Calculator (PIC) web server [78]. The interaction energy of identified residues was evaluated using the amino acid interaction (AAI) web server [79]. PyMOL was used to visualise the resulting interactions. For each protein, prodomain segment interacting with the catalytic domain’s active pocket residues was identified and extracted into a Fasta file. From the interaction energies, residues within these inhibitory segments forming strong contacts with subsite residues were identified. Based on the identified hot spot residues, our next objective was to design short peptide(s) exhibiting the native prodomain effect whilst showing selectivity on human cathepsins. The conservation of prodomain inhibitory segments for all the proteins, and separately of only the plasmodial proteases, was determined using WebLogo server [80]. Peptides of varying lengths and composition based on amino acid conservation forming contacts with subsite residues were proposed. In order to evaluate the interaction of selected peptides on the catalytic domains, the prodomain segments of all proteins were chopped using PyMOL. Blind docking simulation runs of selected peptides were then performed on these sets of catalytic domains by CABS-dock protein-peptide docking tool [81] using the default parameters. To confirm the reliability of the results, docking experiments were repeated using catalytic domains of the same proteins that had been modelled and used in our previous studies [50]. Binding affinity (ΔG) and dissociation constant (Kd) for each protein-peptide complex was then evaluated using PROtein binDIng enerGY prediction (PRODIGY) web server [82].

## Results and discussion

In this work, using combined *in silico* approaches the differences between falcipains and their plasmodial homologs as well as human cathepsins have been evaluated. Based on observed differences and interaction energy profiles between prodomain and catalytic domain subsite residues, short peptides that could mimic the native prodomain inhibitory mechanisms were proposed.

### Both plasmodial and human cathepsins have similar physicochemical properties

Protein function is largely governed by its structure, amino acid composition as well as its environment. Despite the low sequence identity between the two subclasses (cathepsins and plasmodial proteases), physicochemical analysis revealed that they have similar aromaticity and grand average hydropathy (GRAVY) values indicating that both groups of proteins are hydrophilic (Table 3). With an exception of CP-2, all the other proteins have an instability index score of ≤ 40 and thus can be considered as being stable in test-tube environment [83]. Interestingly, there is no significant difference between the aromaticity, GRAVY and instability index scores of partial zymogen complex and individual catalytic domains either. However, significant differences exist in the molecular weight and isoelectric point (pI). Plasmodial partial zymogens have higher molecular weight than that of human cathepsins, as they have longer sequences (two additional structural catalytic domain inserts and longer prodomains). A key factor that controls the functioning of cysteine proteases is pH of the milieu in which they are found. All the plasmodial prodomain-catalytic complexes and Cat-L have a slightly acidic pI of 5.66 ± 0.37 with their catalytic domains exhibiting lower pI. The other cathepsins have basic pI for both their partial zymogen complexes and catalytic domains. This difference in pI profiles might explain the localization aspects of these proteins where the plasmodial proteases and Cat-L are found in acidic food vacuoles and lysosomes respectively while the remaining cathepsins are predominantly found in extracellular matrix.

**Table 3.** A summary of physicochemical properties of FP-2 and FP-3 and homologs partial zymogen sequences. Included also are properties where catalytic domain significantly varied from partial zymogen sequence.

| Protein | Aromaticity | GRAVY | Instability<br>index | Mwgt |  | pI |  |
| --- | --- | --- | --- | --- | --- | --- | --- |
|  |  |  |  | Complex | Catalytic | Complex | Catalytic |
| **FP-2** | 0.12 | -0.59 | 40.31 | 38021.06 | 27176.69 | 6.50 | 4.94 |
| **FP-3** | 0.14 | -0.53 | 33.90 | 38029.91 | 27348.62 | 5.47 | 4.72 |
| **VP-2** | 0.14 | -0.40 | 23.78 | 37941.11 | 27388.93 | 5.49 | 4.74 |
| **VP-3** | 0.14 | -0.47 | 28.92 | 38405.76 | 28088.95 | 5.60 | 5.00 |
| **KP-2** | 0.14 | -0.54 | 29.55 | 38583.83 | 27947.73 | 5.69 | 4.93 |
| **KP-3** | 0.14 | -0.51 | 23.16 | 38284.39 | 27685.36 | 5.89 | 5.13 |
| **BP-2** | 0.14 | -0.52 | 39.73 | 38014.94 | 27367.90 | 5.44 | 4.77 |
| **CP-2** | 0.13 | -0.47 | 52.14 | 37883.92 | 27140.51 | 5.14 | 4.54 |
| **YP-2** | 0.14 | -0.46 | 40.05 | 38120.25 | 27454.06 | 5.66 | 4.78 |
| **Cat-K** | 0.10 | -0.62 | 33.12 | 34566.05 | 23495.47 | 8.83 | 8.92 |
| **Cat-L** | 0.13 | -0.70 | 38.17 | 35074.20 | 24298.86 | 5.33 | 4.64 |
| **Cat-S** | 0.12 | -0.56 | 37.20 | 34986.63 | 23963.97 | 8.44 | 7.64 |

### Plasmodial clan CA proteases and human cathepsins exhibit separate evolutionary clustering

In addition to the previous findings for catalytic domain conservation discussed in detail in ref [7], current MSA identified two highly conserved ERFNIN and GNFD motifs, which are located in the α2-helix and the adjacent downstream loop region between β turn and α3-helix respectively (Figure 1 and Figure 4).

**Figure 4.**
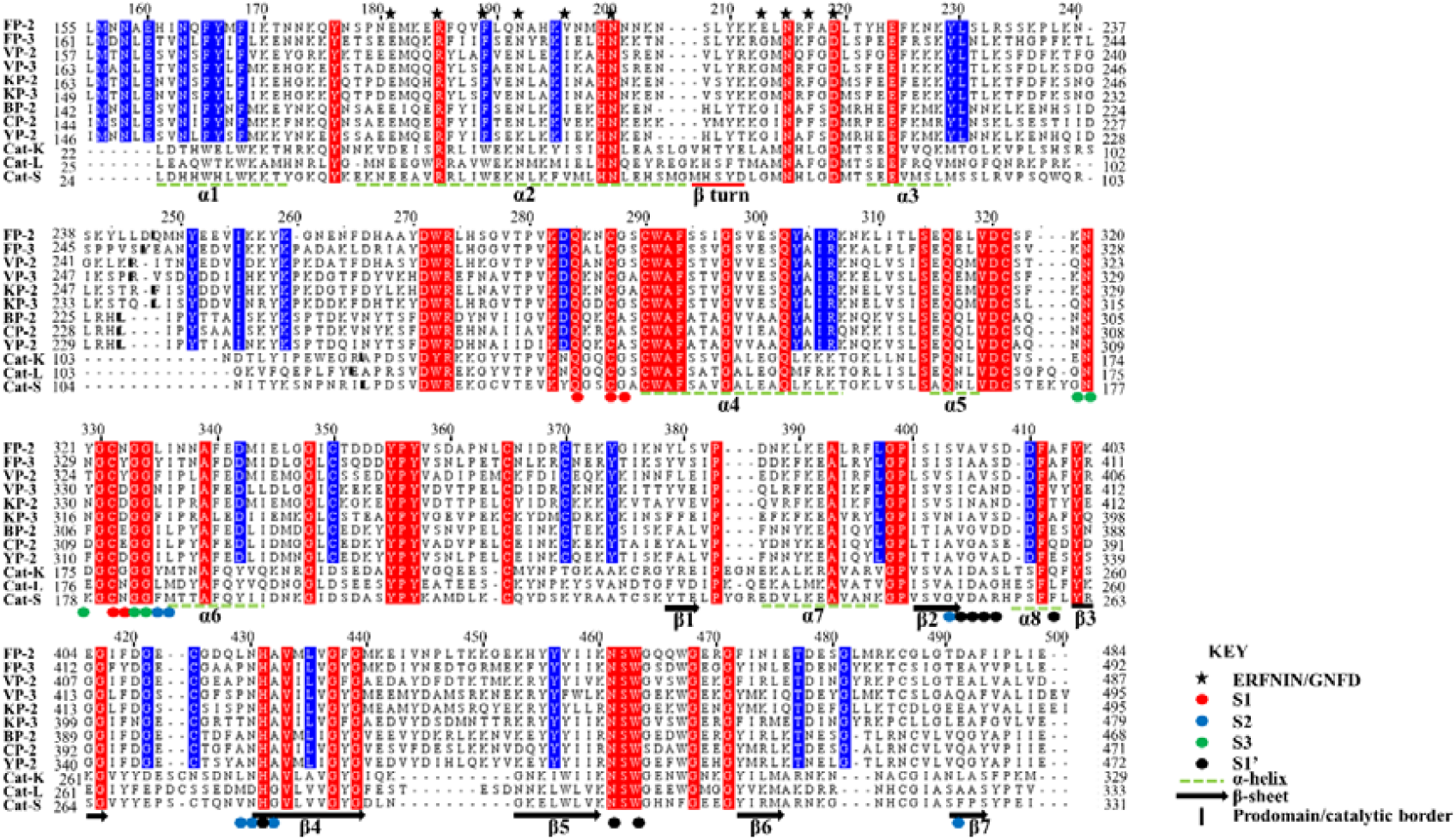
Structural-based multiple sequence alignment of FP-2, FP-3 and homologs prodomain-catalytic domains. Actual residue numbering per protein is given on the side, and the top numbering is based on partial zymogen alignment. The papain family characteristic prodomain ERFNIN and GNFD motif residues are indicated with an asterisk. Bold short lines depict prodomain-catalytic domain border. Dashed green lines indicate the position of α-helix and arrows β-sheet structural elements. Fully conserved residues in all the proteins are marked with red while residues only conserved in plasmodial proteases with blue. Position of subsite residues is shown with filled circles (Red=S1, Blue= S2, Green=S3 and S1’=black).

Despite the highly conserved nature of the ERFNIN motif across all the plasmodial proteins studied, FP-2 and CP-2 have Val residue in the place of Ile196 (numbering based on FP-2). In the human cathepsins, the motif’s Phe190 (FP-2 numbering) is replaced by a Trp, a more hydrophobic residue. Using site-directed mutagenesis, Kreusch *et al*., identified two additional conserved Trp residues in human Cat-L (position 29 and 32 in Cat-L full length protein) which together with the highly conserved motifs (ERFNIN and GNFD) are important in the stability of the partial zymogen complex [84]. In plasmodial proteases, conservative substitution occurs on these two residues whereby they are replaced by less hydrophobic Phe residues (position 165 and 169 in FP-2). The contribution of these amino acid variations will be further discussed in the *“Prodomain regulatory effect mediated by α3 helix hydrophobic interactions with subsites S2 and S1’ residues”* section. MSA result also revealed that cathepsins have a three amino acid insert in the α2 helix between the ERFNIN/GNFD motifs which is absent in the plasmodial proteases, and its importance is yet to be reported.

Phylogenetic analysis using partial zymogen sequences gave a distinct clustering between plasmodial proteins and human cathepsins forming two separate clades (Figure 5). There is no notable difference in tree topology in analysis performed using the catalytic domains only. This can be explained by the observed low sequence identity in both partial zymogen (Table 1) and catalytic domain sequences between the two groups of proteins [50]. The plasmodial proteases further clustered into two main subgroups based on the host. This is attributed to the previously reported sequence variations between the human and rodent plasmodial proteases [50]. FP-2 and FP-3 forms a separate sub-group from the other human plasmodial proteases possibly due to the high sequence similarity between the two proteins. The rate of mutation accumulation appears to vary between the two classes of proteins, being slowest in the human cathepsins. All human plasmodial proteases seem to evolve at the same rate as compared to the rodent orthologs which appear to show the highest substitution rate among all the proteins.

**Figure 5.**
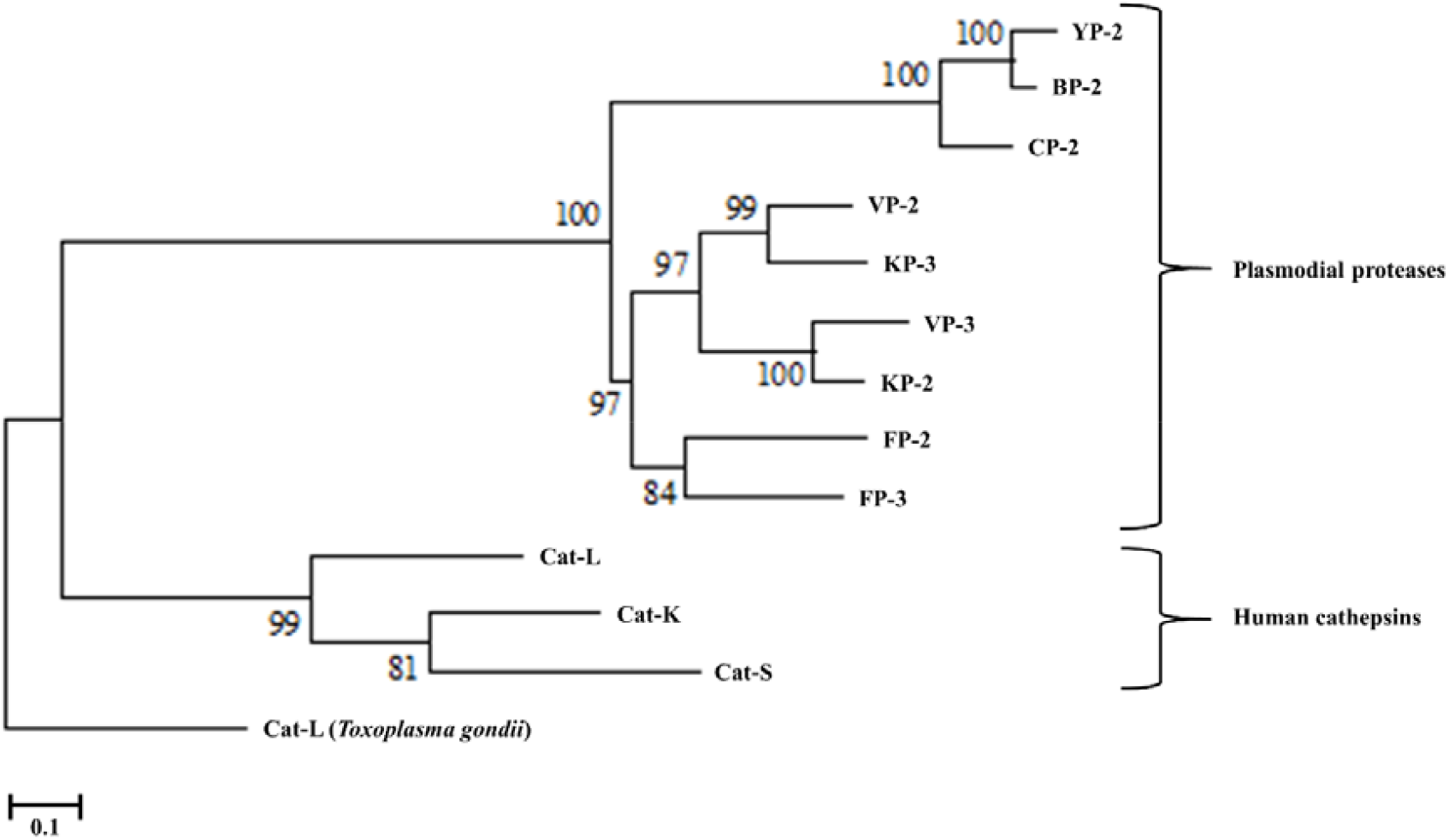
A phylogenetic tree of plasmodial and human FP-3, and FP-3 homologs prodomain-catalytic protein sequences using MEGA5.2.2. The evolutionary history was inferred by using the Maximum Likelihood method based on the Whelan and Goldman model (WAG) model with a γ discrete distribution (+G) parameter of 2.4 and an evolutionary invariable ([+I]) of 0.1. All positions with gaps were completely removed (100% deletion) and bootstrap value set at 1,000. The scale bar represents number of amino acid substitutions per site. *Toxoplasma gondii* CAT-L is used as the outgroup.

### Plasmodial proteases have unique motifs compared to human cathepsins

Sequence motifs within proteins might be associated with a specific biological function. Thus to better understand and characterise a group of proteins, identification of common and distinguished motifs is of critical importance. A total of 13 unique motifs with varied distributions were identified in the set of proteins studied (Figure 6A). These motifs were then mapped onto the 3D structures of partial zymogen complexes (Figure 6B and 6C). Five motifs (M1, M3, M5, M6 and M7) are present in both the plasmodial and human proteases. Out of these five motifs, M1, M3, M5 and M7 are located at the catalytic domain of all proteins while M6 is at α3-helix region of the prodomain (Figure 6B and 6C). Up to three motifs; M2, M4 (located in α1-helix) and M8 (nose region) are only found within the plasmodial proteases, except FP-2 lacks M8. A differential motif composition of the anterior prodomain region (1-3 helix) of the two classes of proteins was observed with one long motif (M4) in plasmodial proteases while human cathepsins have two (M10 and M12).

**Figure 6.**
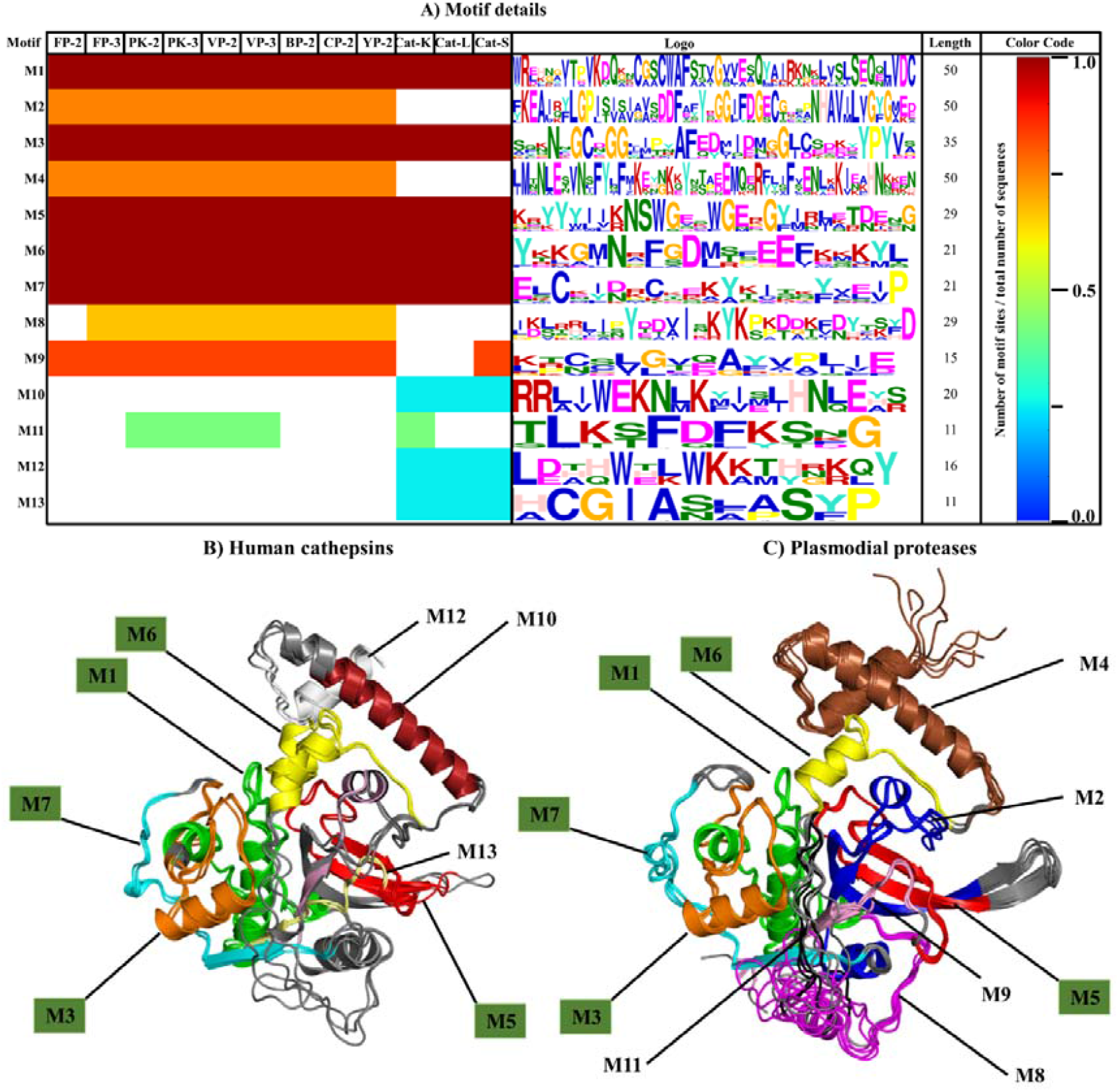
Motif analysis of plasmodial proteases and human cathepsins partial zymogen domains. A) A heat map showing the distribution, level of conservation and information of different motifs found in plasmodial and human proteases studied. A cartoon presentation showing the location of all motifs within the prodomain-catalytic structural fold. Labelled in green boxes are motifs present in both (B) human cathepsins and (C) plasmodial proteases.

PROSITE [85] and MyHits [86] webservers were used to search for the functional importance of identified motifs. M1 (PF00112.15) is the peptidase_C1 functional site and consists of PS00139 (QQnCGSCWAfST-cysteine protease active site), PS00008 (GVvesSQ-N-myristoylation site), and PS00006 (casein kinase II phosphorylation site). M2 (PF00112) is a characteristic functional site of papain-like family cysteine proteases located at the C-terminus (α7-helix to β4), and forms part of the arm region of plasmodial proteases. M3 is located in α6-helix, and the adjacent loop regions of all the Clan CA group of enzymes have no function assigned to it. M4 (PF08246) is known as the cathepsin propeptide inhibitor domain (Inhibitor I29), and is located at α1 and α2 helixes of the N-terminus. The other motifs had no defined function assigned to them according to these webservers.

### Prodomain regulatory effect mediated by α3 helix hydrophobic interactions with subsites S2 and S1‘ residues

Different non-canonical interactions were identified between the prodomain and catalytic domain of proteins. These included hydrophobic, cation-π, ionic, aromatic-aromatic and hydrogen bonds. In all partial zymogen complexes studied, no disulphide linkages between the two domains were observed. The main interactions exhibited are hydrophobic and hydrogen bonds, which participated either in anchoring and maintaining the folding integrity of the prodomain segment, or in mediating its inhibitory effect by interacting with subsite residues (Table S4). Our residue interaction results revealed that prodomain anchoring residues are located on the region between α1-helix and the β-turn which interacted with β3 and part of the arm region in the catalytic domain. Additionally, the C-terminus of the prodomain interacts with the N-terminus of the catalytic domain and part of β3 (Figure 7). A strong hydrogen and hydrophobic interaction network running from the N-terminal end to the GNFD motif prodomain residues, possibly for maintaining its structural fold, was identified in all proteins. In comparison with the human cathepsins, the plasmodial proteases had longer N-terminal prodomain regions with a series of highly conserved residues *viz*. Met156, Asn158, Glu160 and Asn163 (FP-2 numbering). These residues formed a hydrophobic interaction network with bonds of the order < −10.0 kJ/mol. Two additional aromatic-aromatic interactions between Phe165-Phe168 and Phe/Tyr166-Phe189 (FP-2 numbering) in all the plasmodial proteases form strong bonds with energies less than −20.0 kJ/mol and −10.0 kJ/mol respectively. In human cathepsins, the first three Phe positions are substituted with Trp while the fourth position has charged residue substitution (Arg) resulting in weak interactions. A strong residue interaction network between the ERFNIN-GNFD motifs exists in all proteins, confirming the importance of these two motifs in the stability of the prodomain.

**Figure 7.**
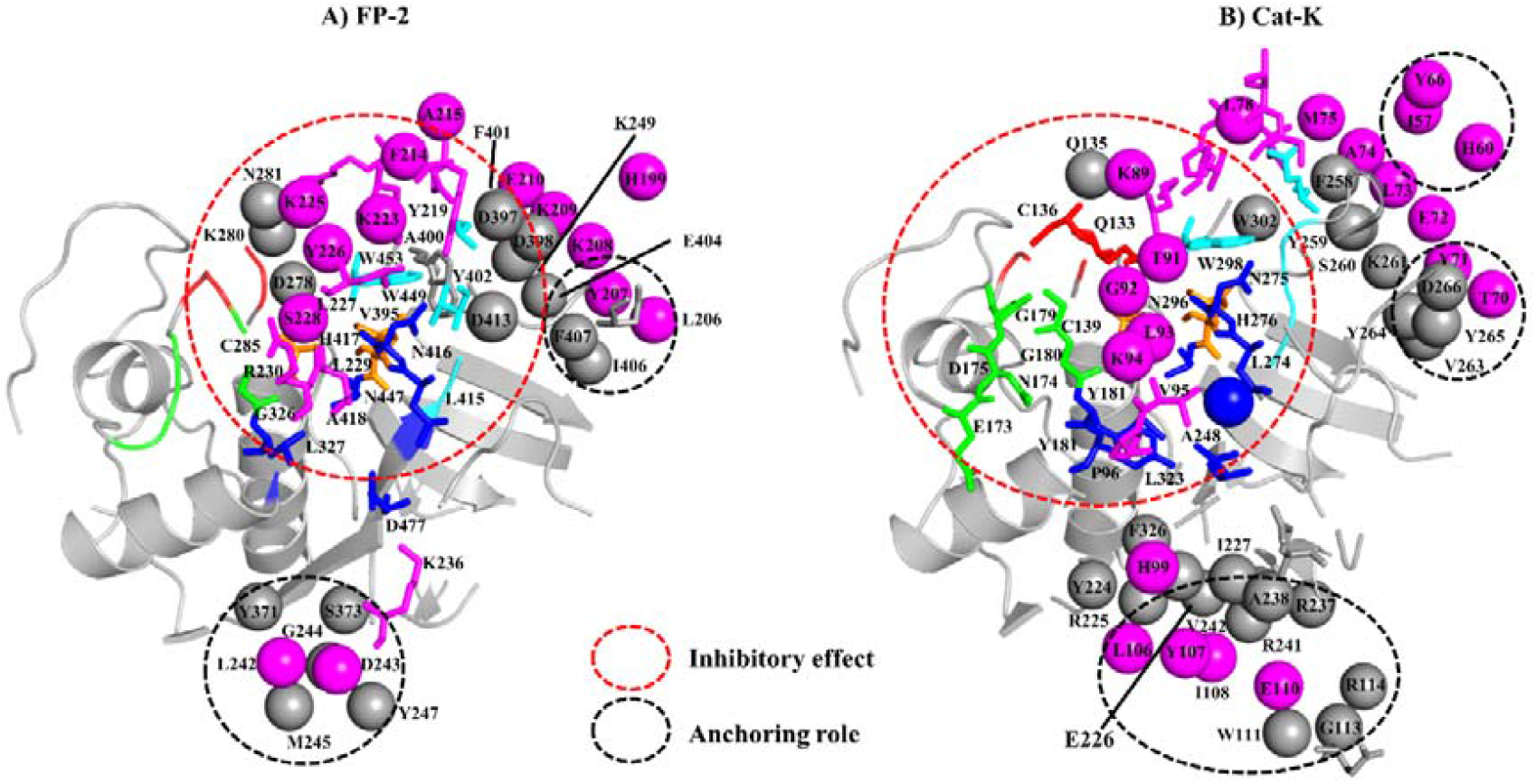
Prodomain-catalytic residue interaction network in (A) FP-2 and (B) Cat-K. For each protein full length residue numbering is used. Prodomain residues (magenta spheres) interacting with non-subsite residues (grey spheres) while sticks are prodomain residues interacting with subsite residues: S1 (red), S2 (blue), S3 (green) and S1’ (cyan). Orange sticks are characteristic catalytic residues of the papain family of proteases (Cysteine, Histidine and Asparagine). Enclosed in red are prodomain-catalytic domain residues mediating inhibitory effect while those in black are involved in anchoring of the prodomain onto the catalytic domain.

A previous mutagenesis study on FP-2 identified two salt bridges (Arg185-Glu221 and Glu210-Lys403) that are important in the activation of the enzyme [87]. In our residue interaction analysis, Arg185 formed a stronger salt bridge with Asp216 (−21.2 kJ/mol) than with Glu220 (−9.5 kJ/mol). To validate these results, Asp216 and Glu220 were independently mutated with an alanine residue and their interaction energy contribution with Arg185 was determined. A complete loss of interaction for Glu70Ala mutation was observed (0.6 kJ/mol) while Asp65Ala energy dropped by half to −12.5 kJ/mol, an indication that the ionic pair between Arg31 and these two positions play a critical biological function. These two residues are fully conserved in all of the proteins studied here. The second predicted salt bridge by Glu210-Lys403 (FP-2 numbering) has high residue variation across the proteins. For the charged Glu210 position in FP-2, all the other plasmodial proteases and Cat-S have a polar residue (Gly) while the other cathepsins have a non-polar residue (Ala). The majority of the residues in position Lys403 are mainly charged except KP-3, CP-2 and Cat-K in which have a polar residue. The energetic contribution from the interactions forming this second salt bridge were all weak (< −1.0 kJ/mol). However, PIC interaction results showed that position 209 in FP-2 consisted of highly conserved positively charged residue (mostly Lysine) across the other plasmodial proteases which formed strong ionic contacts with Asp398 (fully conserved in all plasmodial proteases) in the α8-helix region, an indication that the second salt bridge was most likely formed by these residues. In addition, the mutagenesis study identified aromatic-aromatic interactions in FP-2 between Phe214, Trp449 and Trp453 to be also important in the activation. These residues were conserved in all proteins and formed strong interactions, an indication that they are of functional importance as in FP-2.

A specific aim of this study was to determine the responsible residues that confer the prodomain with its inhibitory function. From residue interaction results, only a small portion of the prodomain (~22-mer) had significant contacts with individual protein subsite residues and was responsible for the inhibitory effect (Figure 8). The main residues mediating the inhibitory effect are located between the α3-helix and the inter-joining loop region, which mostly interact with subsite S2 and S1’ residues via hydrophobic interactions and hydrogen bonds (Table S4). This correlates to our previous findings where residues forming these two subsites were found to be critical in the inhibitory effect and selectivity using non-peptide inhibitors [50].

**Figure 8.**
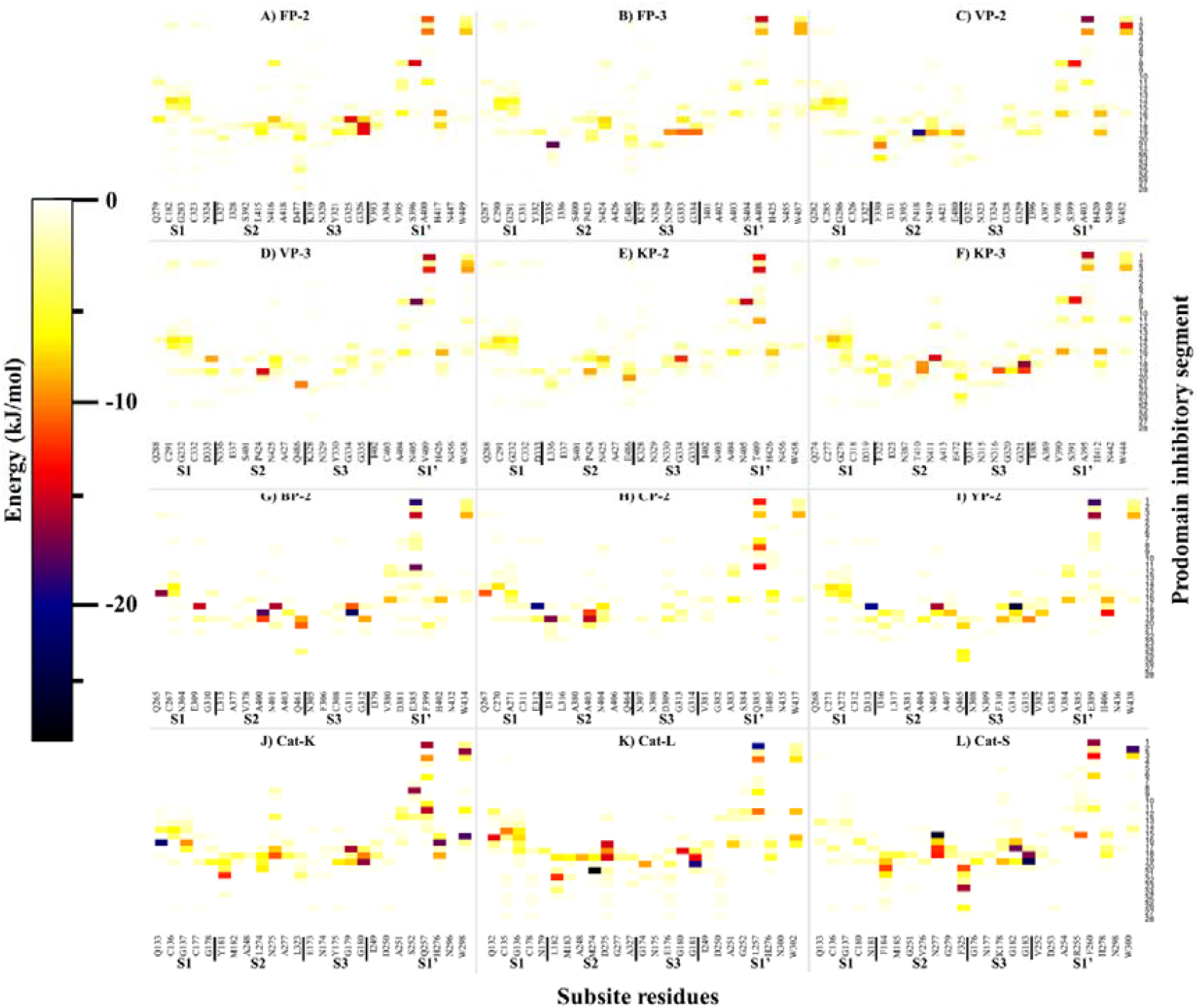
A heatmap for residue interaction energies between prodomain inhibitory segment and the catalytic subsite residues per protein. The inhibitory segment starts from the conserved Asn residue in the GNFD motif (Figure 4).

A common interaction profile between the prodomain inhibitory segment and the catalytic subsites of the different proteins is observed (Figure 8). For subsite S1, a limited residue contact network was observed mainly with residues located at the α3-helix in all the proteases. The C-terminal end of prodomain segment mainly exhibits contacts with S2 and S3 subsites, with human cathepsins and rodent plasmodial proteases forming stronger interactions than the human plasmodial counterparts. In our previous study, high residue variation across the proteins in S2 as well as S1’subsites was reported [50]. In all the proteins, the first three prodomain inhibitory segment amino acids form strong hydrophobic contacts with residues at the opening of S1’ subsite (Figure 8). Rodent plasmodial proteases have additional hydrogen bonding network, due to presence of charged residues at the fourth or fifth S1’ position. From the interaction energy results, there are no observable contacts between residues Ala218 to Thr221 (FP-2 numbering) with any of the proteins’ subsite residues. However, a strong hydrogen bonding network is formed between prodomain Ser231, Leu232, Arg233 with Leu429, Asn430 (S2), Val406, Ala412 (S1’) and Gly334-335 (S3). Lys236 residue forms very strong ionic interactions with Asp491 (S2), a position mainly occupied by charged residues only in the human plasmodial proteases. The side-chain of Ser228 in FP-2 forms hydrogen bonding with thiol group of catalytic Cys285. A similar trend with other plasmodial proteases was observed (Table S4). From the interaction fingerprint, residues that are key in anchoring and maintaining the stability of the prodomain as well as mediating its catalytic domain regulatory effect were identified per protein.

### Peptide inhibitory effect and selectivity dependent on composition and length

Despite their poor chemical properties, peptides remain a promising class of enzyme modulators as they are chemically diverse, highly specific and relatively safe [88,89]. Designing peptide based inhibitors requires prior understanding of how an enzyme recognizes its native peptide substrate then modifying the resulting interactions. Additionally, hot spot residues that regulate protein-protein/domain interactions may provide valuable insights. For FP-2, three peptide studies based on its prodomain-catalytic domain interaction network have already been performed. Rizzi *et al*., who designed peptidomimetics based on the interaction information between cystatin and FP-2 [90]. A major limitation of this study was that it was limited to FP-2 and the broad inhibitory potency of resulting cystatin mimics to other plasmodial proteases was necessary. Another study by Korde *et al*., using a synthetic 15-mer oligopeptide based on the N-terminal extension of FP-2 partial zymogen (LMNNAEHINQFYMFI) showed that it could inhibit substrate processing activity of recombinant FP-2 *in vitro* [91]. However, our interaction fingerprint results showed that this terminal extension was not the native inhibitory segment and was not interacting with any of FP-2 catalytic domain subsite residues. Lastly, Pandey *et al*., expressed the whole prodomain of FP-2 together with truncated segments and evaluated their inhibitory ability against a series of papain-family cysteine proteases. At the end, they determined that a FP-2 prodomain segment (Leu127-Asp243) which included the ERFNIN and GNFD motifs had a broad inhibitory activity against FP-3, BP-2, FP-2, Cat-L, Cat-B and cruzain [92]. Considering its length and molecular mass, the therapeutic potential of this peptide is uncertain.

In our study, peptides aimed at mimicking the inhibitory prodomain segment were designed and tested based on the identified prodomain-catalytic domain interaction fingerprint (Figure 8). Initially, a 22-mer peptide (peptide 1 = NRFGDLSFEEFKKKYLNLKLFD) based on the conservation of the prodomain segments responsible for the inhibitory mechanism for all the proteases was selected for docking against the catalytic domains of individual proteins using the CABS-dock webserver (Figure 9). CABS-dock performs blind docking simulations to identify the most probable binding site while maintaining the flexibility of the peptide ligand [81]. The ΔG of top protein-peptide complex model per protein was then determined using the PRODIGY server. A portion of this peptide interacted with active pocket residues of individual proteins and formed complexes exhibiting high binding affinities as that of a FP-2/Chagasin X-ray crystal complex [PDB: 2OUL] (Table 4). Despite the high predicted affinity scores with peptide 1, no differential binding was observed with the human cathepsins. As its N-terminus had highly conserved GNFD motif residues responsible for anchoring and maintaining the prodomain integrity, we chose to find out if a shorter peptide lacking these residues would bind differently. Thus, a different set of docking experiments with a peptide (peptide 2 = LTYHEFKNKYLSLRSSK) derived from the main inhibitory segment of FP-2 was performed. Despite the variation in length, peptide 2 had similar results to peptide 1 and lacked differential binding affinity profile between the two protein classes.

**Figure 9.**
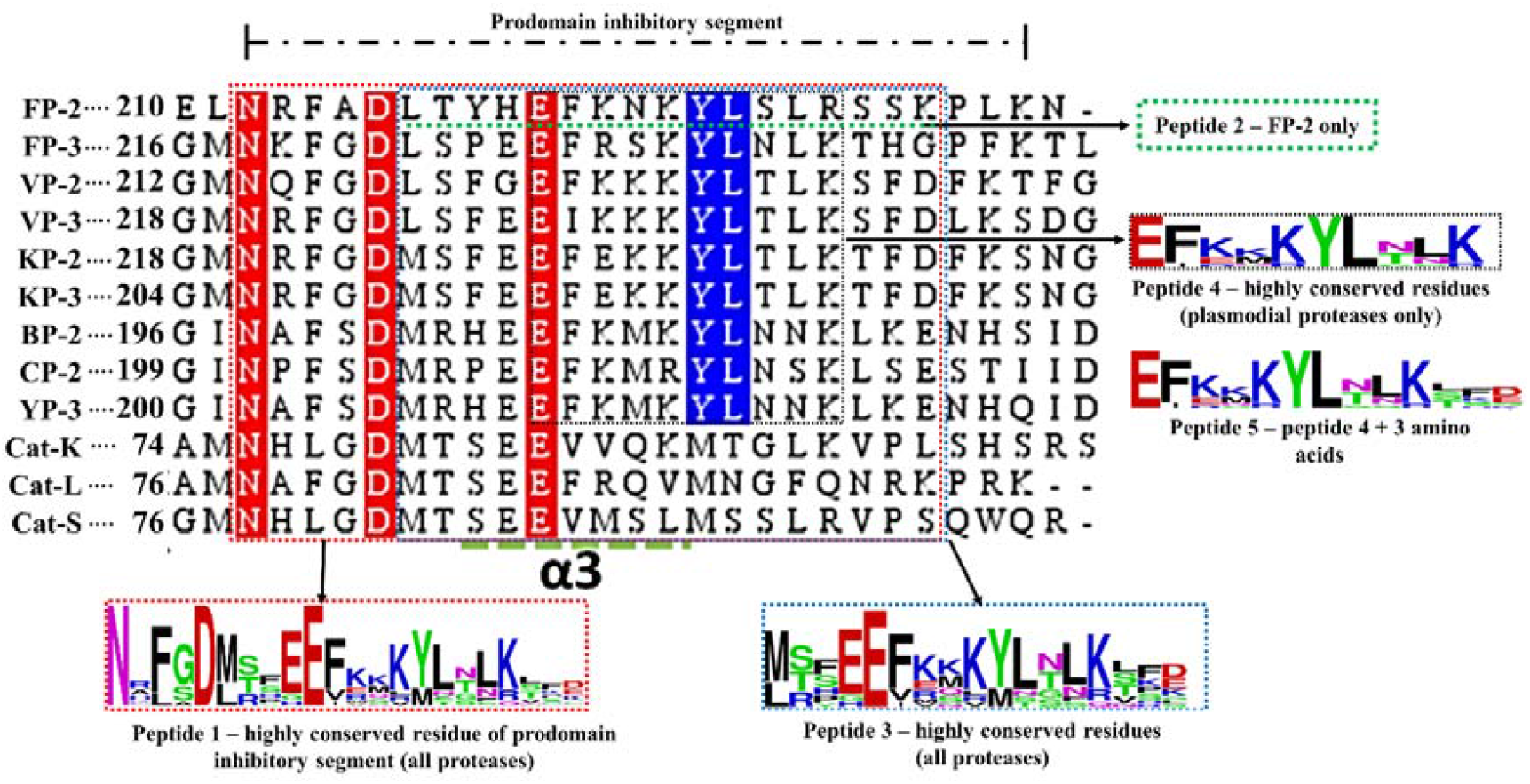
Sequence alignment of the prodomain inhibitory segment for the plasmodial and human cathepsin proteases studied. Marked sequence sections indicate the portions used to design different oligopeptides for docking studies and their conservation as determined by WebLogo server. Actual residue numbering per protein is given on the side.

**Table 4.** Amino acid sequences of proposed peptides, their predicted binding affinity values (ΔG - kcalmol^−1^) and dissociation constant (K_d_) with individual catalytic domains of the different proteins studied.

| Protein | Peptide |  |  |  |  |  |  |  |  |  |
| --- | --- | --- | --- | --- | --- | --- | --- | --- | --- | --- |
|  | 1 |  | 2 |  | 3 |  | 4 |  | 5 |  |
| | $\Delta G$ | $K_D$ (M) | $\Delta G$ | $K_D$ (M) | $\Delta G$ | $K_D$ | $\Delta G$ | $K_D$ | $\Delta G$ | $K_D$ (M) |
| FP-2 | -11.8 | 2.2E-09 | -11.2 | 8.4E-09 | -8.1 | 1.2E-06 | -9 | 2.7E-07 | -11.4 | 4.4E-09 |
| FP-3 | -12.2 | 1.2E-09 | -10.5 | 2.0E-08 | -9.6 | 9.4E-08 | -9.7 | 7.8E-08 | -12.3 | 8.8E-10 |
| VP-2 | -11.0 | 8.3E-09 | -10.1 | 3.8E-08 | -12.3 | 8.8E-10 | -9.0 | 2.7E-07 | -10.9 | 1.1E-08 |
| VP-3 | -11.7 | 2.7E-09 | -11.0 | 8.3E-09 | -9.9 | 5.7E-08 | -9.2 | 1.7E-07 | -11.8 | 2.1E-09 |
| KP-2 | -12.7 | 4.7E-10 | -11.8 | 2.1E-09 | -6.7 | 1.3E-05 | -8.1 | 1.2E-06 | -10.6 | 1.8E-08 |
| KP-3 | -11.8 | 2.1E-09 | -12.7 | 5.2E-10 | -9.3 | 1.4E-07 | -9.2 | 1.9E-07 | -12.7 | 5.2E-10 |
| BP-2 | -11.5 | 3.7E-09 | -12.2 | 1.2E-09 | -10.9 | 9.9E-09 | -7.9 | 1.7E-06 | -11.7 | 2.7E-09 |
| CP-2 | -11.9 | 2.2E-09 | -11.7 | 2.7E-09 | -8.7 | 3.9E-07 | -8.3 | 1.1E-07 | -12.7 | 4.7E-10 |
| YP-2 | -11.7 | 2.7E-09 | -14.3 | 5.5E-11 | -12.4 | 7.9E-10 | -10.4 | 2.5E-08 | -9.3 | 1.5E-07 |
| Cat-K | -13.9 | 6.3E-11 | -11.5 | 3.5E-09 | -11.9 | 2.0E-09 | -11.4 | 4.2E-09 | -8.3 | 1.1E-07 |
| Cat-L | -12.2 | 1.0E-09 | -11.8 | 5.1E-09 | -12.3 | 9.7E-10 | -10.2 | 3.5E-08 | -8.7 | 9.2E-07 |
| Cat-S | -10.4 | 2.4E-09 | -11.5 | 3.5E-09 | -12.4 | 7.7E-10 | -9.9 | 5.2E-08 | -8.6 | 4.8E-07 |

\*FP-2 complex with chagasin: $\Delta G$ -11.9, $K_D$ =1.9e-09.
Peptide 1<sup>NRF</sup>GDLSFEFFKKKYLNLKLF<sup>D</sup>, 2<sup>LTY</sup>HEFNKYL<sup>SL</sup>RSSK, 3<sup>MT</sup>FEEFKQKYL<sup>TLK</sup>SKD, 4<sup>EF</sup>KNKYL<sup>TLK</sup>, 5<sup>EF</sup>KNKYL<sup>TLK</sup>SKD

A previous *in vitro* study by Pandey *et al*., show that a FP-2 prodomain harbouring peptide 2 segment exhibited similar broad inhibitory activity on cruzain, Cat-B, Cat-L, FP-2, FP-3 and BP-2 [92]. However, from our energy interaction profiles, a large portion of the tested prodomain including the ERFNIN/GNFD motifs is mainly involved in anchoring it to the catalytic domain. Thus, the main inhibitory segment is much shorter and downstream of the GNFD motif. Peptide 2-YP-2 complex had the strongest binding association (−14.3 kcal/mol) while VP-2 had the lowest (−10.1 kcal/mol). With the already tested peptides exhibiting unselective high affinity binding on both human cathepsins and plasmodial proteases, additional docking experiments were performed with a different peptide derived from the most conserved residues in the same inhibitory segment as peptide2 from all proteases (peptide 3 = MTFEEFKQKYLTLKSKD). In some positions within the prodomain inhibitory segments across the plasmodial proteases, high residue variations were observed and there was no consensus about which residue to include in the peptide. So the properties of the residues (polar, charged, non-polar, hydrophobic) occupying such positions were compared to determine if a common chemical property was preferred.. In addition, residues showing stronger interactions with the catalytic subsite residues were also taken into account. However, the ΔG between peptide 3 and plasmodial proteases was significantly lower in most plasmodial proteases than with the earlier peptides. Nevertheless, human cathepsins had similar binding affinity values with peptide 1 and 2. Guided by the residue interaction profile of prodomain residues with subsite residues (Figure 8 and S2), a fourth peptide (peptide 4 = EFKNKYLTLK) composed of the most conserved amino acids around α3-helix of the inhibitory segment of all plasmodial proteases was evaluated. A similar trend of nonselectivity was observed as with peptide 1, though with lower binding affinity. A fifth peptide, similar to peptide 4 except for its length, (EFKNKYLTLKSKD) was also evaluated. The residues in this peptide showed some conservation in the plasmodial proteases and had significant differences to the human cathepsins. Interestingly, it bound more strongly to all plasmodial proteases compared to the human cathepsins. A likely explanation of this differential binding affinity was that the peptide interacted with fewer residues on human cathepsins compared to the plasmodial proteins (Figure 10). In most of the plasmodial proteases, peptide 5 bound with almost same affinity as that of chagasin and FP-2 (−11.9 kcal/mol). From the prodomain-catalytic interaction analysis (Table 5 and Figure 10), the terminal end in peptide 5 interacts with last position of S2 which consists of a charged residue (only in human plasmodial proteases) forming a strong ionic interaction as well as other nonsubsite residues thus forming a stronger complex. In most plasmodial proteases, peptide 5 formed multiple hydrogen bonds especially with S2 and S1’ subsite residues. These two subsites residues have been found to be key in determining binding selectivity as they are the main contributors to ligand binding [50]. Docking studies with previously modelled catalytic domains gave results consistent with the current models.

**Figure 10.**
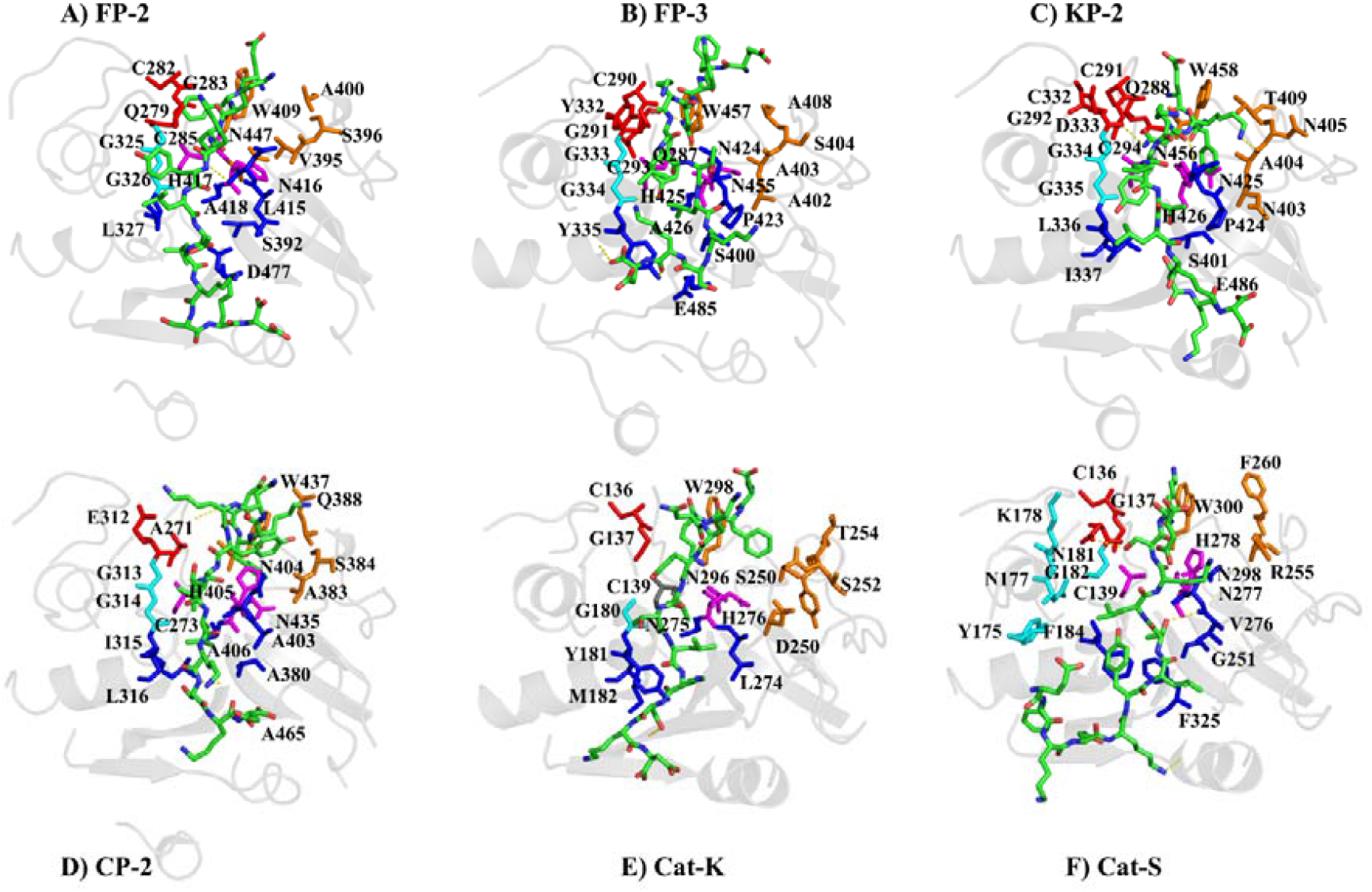
Peptide 5-catalytic domain subsite residue interactions of various proteins (Red=S1, Blue=S2, Cyan=S3, S1’=orange and Magenta=catalytic residues).

**Table 5.**
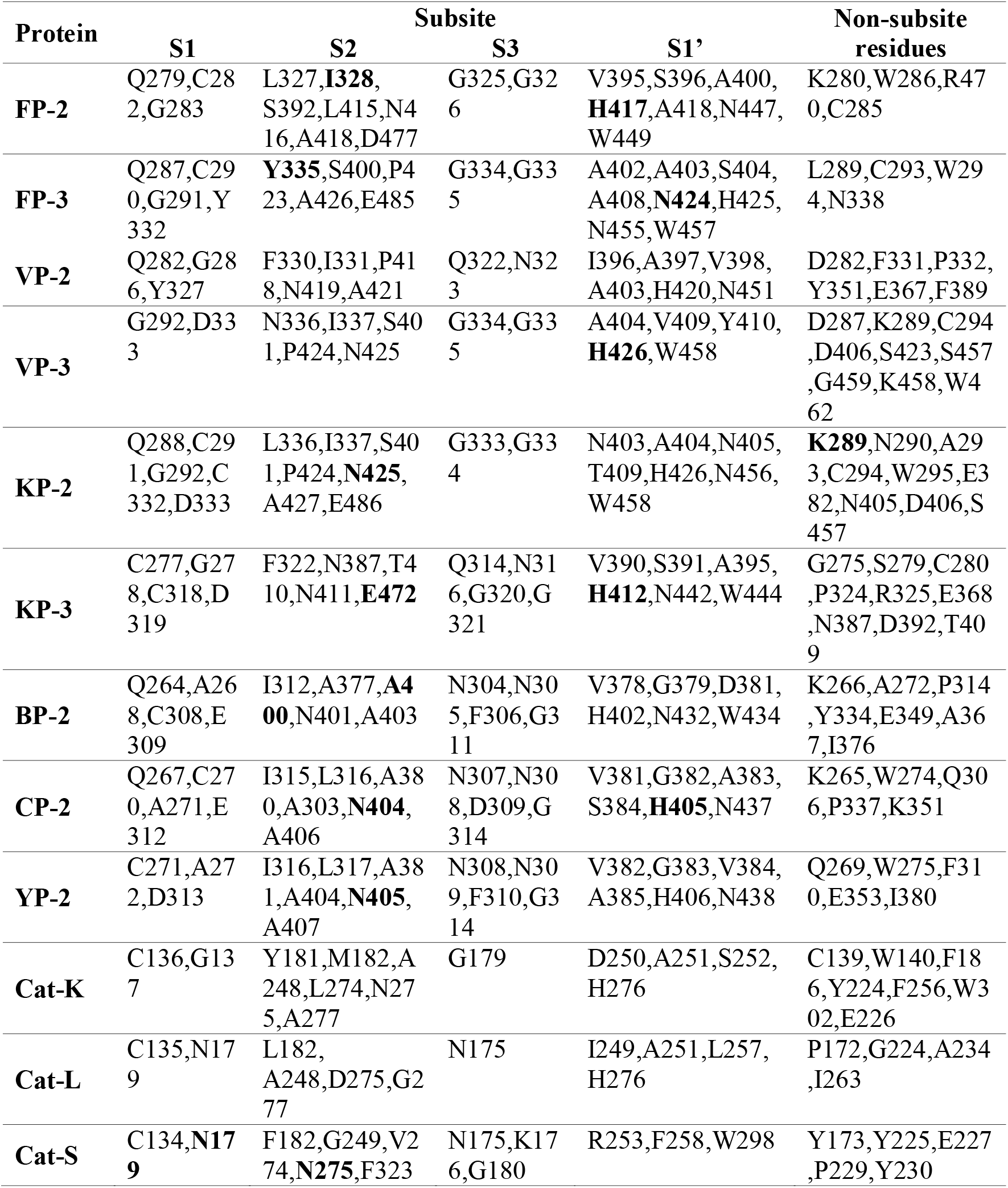
Peptide 5-catalytic domain residue interaction fingerprint. Bold are residues forming hydrogen bonds with the peptide.

| Protein | Subsite |  |  |  | Non-subsite residues |
| --- | --- | --- | --- | --- | --- |
|  | S1 | S2 | S3 | S1' |  |
| **FP-2** | Q279,C28<br>2,G283 | L327,**I328**,<br>S392,L415,N4<br>16,A418,D477 | G325,G32<br>6 | V395,S396,A400,<br>**H417**,A418,N447,<br>W449 | K280,W286,R47<br>0,C285 |
| **FP-3** | Q287,C29<br>0,G291,Y<br>332 | **Y335**,S400,P4<br>23,A426,E485 | G334,G33<br>5 | A402,A403,S404,<br>A408,**N424**,H425,<br>N455,W457 | L289,C293,W29<br>4,N338 |
| **VP-2** | Q282,G28<br>6,Y327 | F330,I331,P41<br>8,N419,A421 | Q322,N32<br>3 | I396,A397,V398,<br>A403,H420,N451 | D282,F331,P332,<br>Y351,E367,F389 |
| **VP-3** | G292,D33<br>3 | N336,I337,S40<br>1,P424,N425 | G334,G33<br>5 | A404,V409,Y410,<br>**H426**,W458 | D287,K289,C294<br>,D406,S423,S457<br>,G459,K458,W4<br>62 |
| **KP-2** | Q288,C29<br>1,G292,C<br>332,D333 | L336,I337,S40<br>1,P424,**N425**,<br>A427,E486 | G333,G33<br>4 | N403,A404,N405,<br>T409,H426,N456,<br>W458 | **K289**,N290,A29<br>3,C294,W295,E3<br>82,N405,D406,S<br>457 |
| **KP-3** | C277,G27<br>8,C318,D<br>319 | F322,N387,T4<br>10,N411,**E472** | Q314,N31<br>6,G320,G<br>321 | V390,S391,A395,<br>**H412**,N442,W444 | G275,S279,C280<br>,P324,R325,E368<br>,N387,D392,T40<br>9 |
| **BP-2** | Q264,A26<br>8,C308,E<br>309 | I312,A377,**A4**<br>**00**,N401,A403 | N304,N30<br>5,F306,G3<br>11 | V378,G379,D381,<br>H402,N432,W434 | K266,A272,P314<br>,Y334,E349,A36<br>7,I376 |
| **CP-2** | Q267,C27<br>0,A271,E<br>312 | I315,L316,A38<br>0,A303,**N404**,<br>A406 | N307,N30<br>8,D309,G<br>314 | V381,G382,A383,<br>S384,**H405**,N437 | K265,W274,Q30<br>6,P337,K351 |
| **YP-2** | C271,A27<br>2,D313 | I316,L317,A38<br>1,A404,**N405**,<br>A407 | N308,N30<br>9,F310,G3<br>14 | V382,G383,V384,<br>A385,H406,N438 | Q269,W275,F31<br>0,E353,I380 |
| **Cat-K** | C136,G13<br>7 | Y181,M182,A<br>248,L274,N27<br>5,A277 | G179 | D250,A251,S252,<br>H276 | C139,W140,F18<br>6,Y224,F256,W3<br>02,E226 |
| **Cat-L** | C135,N17<br>9 | L182,<br>A248,D275,G2<br>77 | N175 | I249,A251,L257,<br>H276 | P172,G224,A234<br>,I263 |
| **Cat-S** | C134,**N17**<br>**9** | F182,G249,V2<br>74,**N275**,F323 | N175,K17<br>6,G180 | R253,F258,W298 | Y173,Y225,E227<br>,P229,Y230 |

From the motif analysis (Figure 6), a large proportion of peptide 5 was represented in motif M6. Despite the functional annotation of motif M4 indicating it as the cathepsin propeptide inhibitor domain, majority of its residues were predominantly involved in anchoring the prodomain. Taken together, our study is the first to identify the most key prodomain segment involved in regulation of cysteine proteases, and to apply information based approaches to propose a peptide with differential binding on both human and plasmodial proteases.

## Conclusion

In the present study we aimed to characterise the differences between *P. falciparum* falcipains and their plasmodial and human homologs, especially where the prodomain interacts with the catalytic domain, in order to identify key residues which could be useful in antimalarial drug development approaches. This was done at both sequence and structure level. Through homology modelling, near native 3D partial zymogen complexes of both plasmodial and human proteases were obtained. This allowed structural characterization, thus deciphering how these segments confer their inhibitory mechanism endogenously. The main prodomain residues mediating the inhibitory effect were located in the α3-helix and the inter-joining loop region, and mostly interacted with subsite S2 and S1’ residues. In our previous studies [50,57], we showed that residues forming these two subsites are critical in inhibitor design as they differ from human cathepsins. Hence, putting all the analysis together, with a continuous prodomain epitope mimicking strategy, a peptide which bound selectively more strongly on plasmodial proteases than the human ones was designed. The present approach offers a starting point which could lead to the establishment of novel antimalarial peptide drugs aimed at mimicking the natural plasmodial protease regulatory mechanism. Additional chemical modification either to obtain peptide derivatives with better physicochemical and pharmacokinetic properties as well as potency, bioavailability and stability might be necessary. Accessibility of parasite infected erythrocytes by macromolecules remains a major concern for the development of peptide based antimalarial inhibitors. A study by Farias *et al*., using fluorescent peptides revealed that peptides with molecular weight up to 3146 Da can permeate into the blood stage parasites [93]. All the peptides determined had a mass of below 2753 Da, with peptide 5 having 1613 Da, an indication that it would readily be available inside the parasites. Korde *et al*., demonstrated that a synthetic 15-mer oligopeptide of mass 1885 Da could localise into the intracellular compartments of trophozoites and schizoints inhibiting FP-2 activity [91]. Additional modification of the peptide backbone as well as amino acid side chains may also be performed yielding peptide based inhibitors.

## Acknowledgement

This work is supported by the National Research Foundation (NRF) South Africa (Grant Numbers 93690 and 105267). T.M.M and J.N.N thank Rhodes University for the postgraduate financial support. The content of this publication is solely the responsibility of the authors and does not necessarily represent the official views of the funders.

## Author contributions

Ö.T.B conceived the project. T.M.M and J.N performed the experiments. All authors analysed the data. T.M.M and Ö.T.B wrote the article.

## Abbreviations

Å: Angstrom
AAI: Amino acid interaction
BIC: Bayesian Information Criterion
BP-2: Berghepain 2
CP-2: Chabaupain 2
Cat-K: Cathepsin K
Cat-L: Cathepsin L
Cat-S: Cathepsin S
FPs: Falcipains
FP-2: Falcipain 2
FP-3: Falcipain 3
GRAVY: Grand average of hydropathy index
K_d_: Dissociation constant
KP-2: Knowlesipain 2
KP-3: Knowlesipain 3
MAST: Motif Alignment Search Tool
MEGA: Molecular Evolutionary Genetic Analysis
MEME: Multiple Em for Motif Elucidation
Mr: Molecular weight
MSA: Multiple sequence alignment
NCBI: National Center for Biotechnology Information
NNI: Nearest-Neighbor-Interchange
PIC: Protein Interaction Calculator
pI: Isoelectric point
PlasmoDB: Plasmodium genome Database
PRODIGY: PROtein binDIng enerGY prediction
PROMALS3D: PROfile Multiple Alignment with predicted Local structures and 3D constraints
PSI-BLAST: Position-Specific Iterative Basic Local Alignment Search Tool
RBC: Red Blood Cell
VP-2: Vivapain 2
VP-3: Vivapain 3
YP-2: Yoelipain 2
Z-DOPE: Normalized Discrete Optimized Protein Energy
ΔG: Binding affinity
3D: Three dimensional

